# Batch correction for large-scale mass spectrometry imaging experiments

**DOI:** 10.64898/2026.01.30.702769

**Authors:** Andreas A. Thomsen, Ole N. Jensen

## Abstract

We assess batch correction methods for MALDI mass spectrometry imaging experiments. ComBAT reduced batch-related technical variance, maintained biological variation, and improved the overall score by 19.4%.

**Availability and implementation:** Methods are available in R. comBAT is used through the “sva” package while Harmony, CCA, FastMNN are available in the “Seurat” package https://github.com/satijalab/seurat. scVI, scANVI are Scanorama are available in the Python programming language and through github: https://github.com/scverse/scvi-tools.

## Introduction

Batch effects refer to systematic, non-biological and undesirable variations in data due to differences in experimental conditions across batches [1]. Batch effects occur in all biological analysis regimen and are difficult to avoid, particularly in the context of large-scale omics data, e.g. single cell RNA sequencing (scRNA seq) data [2]. Mass spectrometry imaging (MSI) is an analytical technique that acquires spatially resolved mass spectra of biomolecules (e.g., peptides, metabolites) in a pixel-by-pixel manner by rastering across a specimen, for example, a thin tissue section. A pixel at position (x,y) in the specimen generates a mass spectrum where molecule ions are detected by their mass-to-charge ratio and intensity (m/z, I). The mass spectrum represents the molecular composition at position (x,y). Thus, the data structure in MSI is multidimensional with two spatial dimensions (x,y) and one spectral dimension (m/z,I) per pixel. MSI data can be considered large-scale data in a data frame that typically consists of >100.000 data points (pixels). An experiment may cover 10’s to 100’s of samples and concomitant data acquisition cycles. Thus, batch effects and variations are inherent to MSI [3] due to preanalytical factors including sample preparation protocols, laboratory conditions, data acquisition methods and operators. For example, accuracy of cryotome sectioning of tissue for MSI is sensitive to the tissue structure and texture, leading to variations of tissue thickness. Cumulative batch effects can mask the underlying biological information and may lead to irreproducible or inconclusive results.

We hypothesized that batch correction methods that are commonly used in the omics fields would be applicable to MSI experiments to reduce batch effects while retaining biological variation. We tested the batch correction methods comBAT, Harmony, Seurat canonical correction analysis, and fast mutual nearest neighbor on a large-scale MSI dataset obtained from 128 renal cell carcinoma patients across 26 defined batches in the tissue microarray (TMA) format[4]. We demonstrate successful batch correction using uniform manifold approximation and projection (UMAP) and metrics that scored successful batch integration and conservation of biological variation.

## Material and methods

The clinical samples and MSI experimental setup are described elsewhere (MedRxiv, **doi:** https://doi.org/10.64898/2026.01.13.26343935). MSI experiments were designed to profile trypsin-generated peptides from proteins in the tumor tissue sections. Briefly, the dataset consists of MSI data obtained from 541 needle cores (2- or 3-mm diameter) from 128 renal cell carcinoma (RCC) patients and controls. A total of 26 individual 3 µm tissue microarrays (TMAs, FFPE sections) were deposited onto IntelliSlides (Bruker Daltonics) to enable high-throughput MSI data acquisition from all 541 cores within 48 hours. Thus, the MSI data across 541 cores was acquired in 26 experiments, each consisting of MSI measurements of up to 30 cores. Each RCC core consisted of at least 80% tumor. MSI data (peptide profiles,10 ppm mass accuracy) of 541 cores (26 MSI experiments) were merged into one.imzml file in SCiLS lab v 2023a and loaded into R. The Cardinal package [5] was used for data preprocessing, including total ion count (TIC) normalization, peak picking with signal-to-background ratio of 3, peak alignment with a tolerance of 0.5 Da, peak filtering, peak binning and deisotoping with a tolerance of 6 ppm, and a Pearson correlation coefficient threshold of 0.85. This resulted in a dataset consisting of 866 *m/z* values. Intensity data of the *m/z* values was then Log2-transformed and scaled (centering by mean subtraction from each observation following division by standard deviation) before batch correction. Batch correction methods included comBAT [6], Harmony [7], canonical correction analysis (CCA) from Seurat [8] and fast mutual nearest neighbors (fastMNN) [9]. The latter three methods are integrated and available in the Seurat v5 environment[10], while comBAT is available through the “sva” R package.

In analogy with single cell transcriptomics, we treated individual pixels in MSI as single cells, meaning that one pixel entry {(x,y), (m/z, I)} was considered as one cell for the batch correction algorithm. This is valid as the foundational assumptions of the applied methods are made for imaging-based data and not for characteristics of transcriptomics data (e.g., sparse matrices)[11]. Batches were defined at the tissue microarray section level, resulting in 26 batches, each corresponding to one MSI data acquisition cycle across one TMA section with multiple RCC cores. Harmony, CCA and fastMNN work on dimensionally reduced data that required a principal component analysis (PCA) matrix as input for the correction tool, while comBAT works on the full data frame. The data was subjected to uniform manifold approximation and projection (UMAP) [12] with labelling of batches and conditions before and after correction. This served to assess whether the batch correction methods integrated the batches without influencing the underlying biological variation of the dataset. Additional metrics to assess the efficiency of the batch correction methods included adjusted rand index (ARI)[13], normalized mutual information (NMI)[14], average silhouette width (ASW)[15], PCA regression[16] and Local Inverse Simpson’s Index (LISI)[7] were used. Briefly, ARI and NMI assess biological variation by comparison of cluster assignments against ground truth. ASW considers cluster overlap and separation and depending on the use of labels (batch labels or condition labels) it assesses both data integration and biological variation.LISI scores diversity by measuring the effective number of unique labels within a local neighbourhood. Depending on the label input LISI also assesses both batch mixing and biological variation. PCA regression works by linear regression of the batch variable onto each component, summarizing it for all components and assessing how well the batches are mixed. All metrics were scaled to the interval 0 – 1, where 1 indicates highest batch mixing or conservation of biology variation similarly to Luecken et al [17].

## Results

UMAP analysis of the experimental MSI dataset of 26 batches (MedRxiv, **doi:** https://doi.org/10.64898/2026.01.13.26343935) exhibits two major pixel populations or clusters (Figure 1a). A small cluster contains MSI pixels originating from batches 17 and 26 and a larger cluster contains MSI pixels from the remaining 24 batches. The smaller cluster formed by batches 17 and 26 group the conditions independently from the larger cluster with the remaining batches, which indicates the variation is technical and not biological (Figure 1b). Batch correction by comBAT results in one major homogenous cluster that encompasses all batches (Figure 1c), and the conditions group based on biological-rather than technical variation (Figure 1d).

**FIGURE 1:**
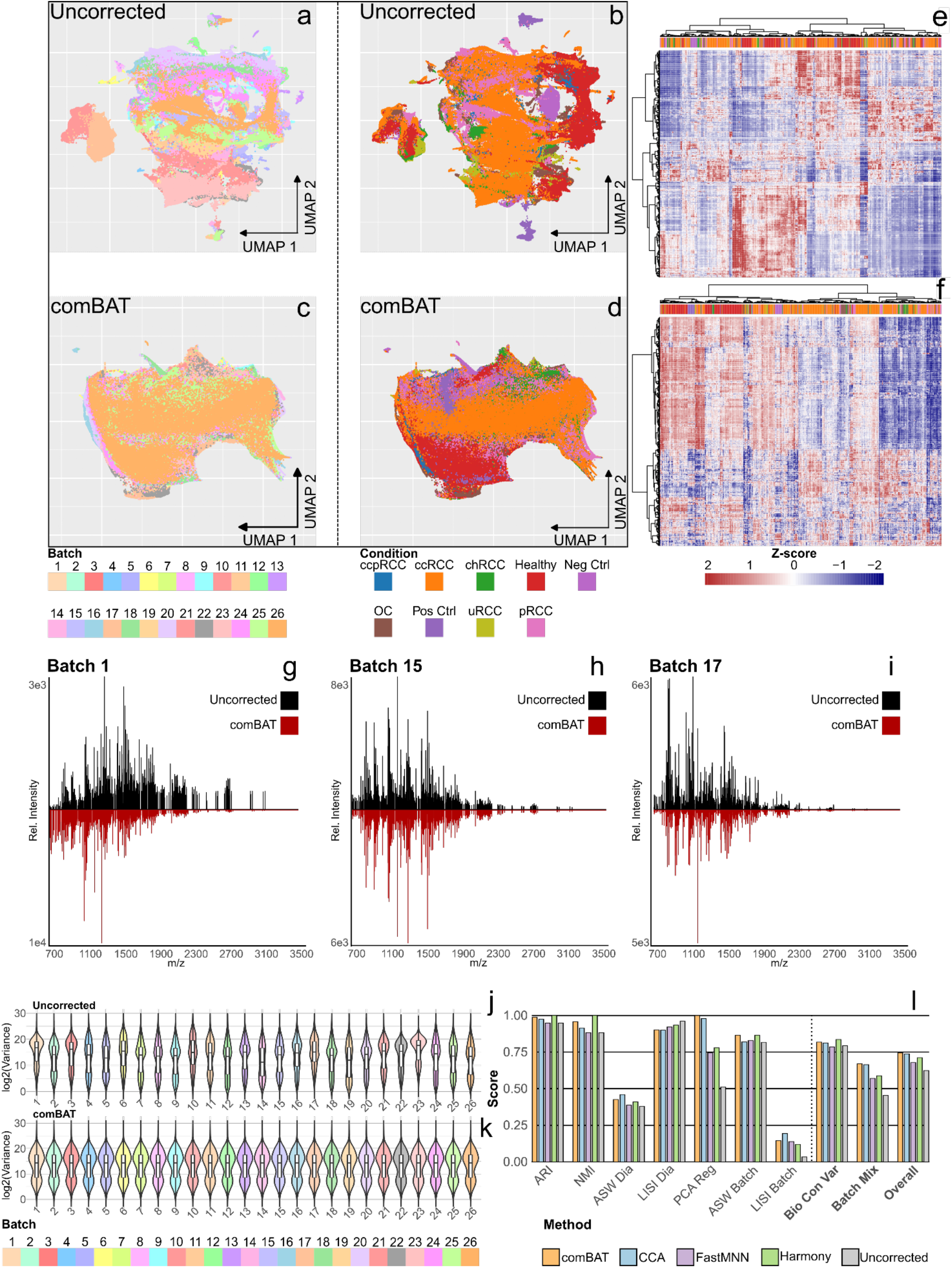
Assessment of batch correction methods for mass spectrometry imaging data. (a) UMAP of uncorrected data, including all pixels in the dataset colored by batch number of origin. (b) UMAP of uncorrected data, including all pixels in the dataset colored by biological origin. (c) UMAP of comBAT corrected data, including all pixels in the dataset colored by their batch number of origin. (d) UMAP of comBAT corrected data, including all pixels in the dataset colored by the biological origin. (e) Heatmap (Z-scores, uncorrected data) by hierarchical clustering of MSI peptide features on the y-axis and hierarchical clustering of tumor subtype cores on the x-axis. (f) Heatmap (Z-scores, comBAT corrected data) by hierarchical clustering of MSI peptide features on the y-axis and hierarchical clustering of tumor subtype cores on the x-axis (g) Extracted mass spectrum before (black) and after comBAT correction (red) for batch 1. x-axis depicts *m/z* and y-axis shows relative intensity. (h) Extracted mass spectrum before (black) and after comBAT correction (red) for batch 15. x-axis depicts *m/z* and y-axis shows relative intensity. (i) Extracted mass spectrum before (black) and after comBAT correction (red) for batch 17. x-axis depicts *m/z* and y-axis shows relative intensity. (j) Violin plots of data variance (log2) of uncorrected MSI data for the 26 batches. x-axis shows the batch number and y-axis depicts log2-scaled variance values. (k) Violin plots of data variance (log2) of comBAT corrected MSI data for the 26 batches. x-axis shows the batch number and y-axis depicts log2-scaled variance values. (l) Assessment of four batch correction methods: comBAT, CCA, FastMNN, Harmony versus uncorrected data. Barplots of metrics used to assess batch mixing and biological variation conservation. ASW Dia, LISI Dia, ARI and NMI assess conservation of biological variation using predefined sample labels. ASW Batch, LISI Batch and PCA reg assess batch mixing using predefined batch labels. Batch Mix is the mean of batch mixing metrics: ASW Batch, LISI Batch and PCA reg. Bio Con Var is the mean of biological variation conservation metrics: ASW Dia, LISI Dia, ARI and NMI. Overall is the mean of all the metrics. x-axis is the methods and y-axis is the score values are in the range of 0 - 1, with 1 indicating better batch mixing or bio variation conservation.

Next, we performed hierarchical clustering analysis before and after batch correction to assess if clustering would improve upon correction with emphasis on conditions. The clustering of uncorrected data (Figure 1e) does not reflect the underlying biology (i.e. the data is classified randomly). After comBAT correction (Figure 1f) the clustering of the samples improves as the experimental conditions healthy, chRCC and OC now cluster together within their class.

We speculated that batch effects originate from size-heterogeneity of the tryptic peptides, as *in situ* digestion kinetics are difficult to control. Short and long tryptic peptides would be more prone to this due to lower abundance compared to peptides in the optimal range of *m/z* 1000-2500. Indeed, the corrected extracted mass spectra exhibited reduced relative peak intensity of long peptides (batch 1, 15 and 17) (Figure 1g-1i) and short peptides (batch 15 and 17) (Figure 1h and 1i), indicating that they vary the most across experimental batches.

Batch correction should result in a reduction of the variance across all batches. This is demonstrated by data shown in Figure 1j where outlier batches 1, 17, 23 and 26 are seen before correction based on their median, density distribution and interquartile ranges (IQRs). After comBAT correction (Figure 1k), the density distributions are similar for all batches together with the median and IQRs, which indicates that some data variation is corrected.

We summarized the batch correction assessment metrics in Figure 1l. Harmony scores highest in terms of conservation of biological variation with a score of 0.84 while comBAT performs best for the batch mixing task with a score of 0.67. All batch correction methods reduced the batch effects and had higher batch mixing scores compared to uncorrected data. All tested methods retained the biological variation of the data with improvements in biological variation conservation scores.Overall, comBAT and CCA correction scored highest for performance (0.74) vs the uncorrected data (0.62) (Figure 1l), i.e. an improvement of 19.4%.

## Discussion

We show that batch correction methods commonly used in the omics fields are applicable to large-scale mass spectrometry imaging experiments. All the tested batch correction methods reduced the batch effects of the MSI dataset (Figure 1e).

Batch correction by comBAT reduced the batch effects across 26 batches of MSI data from RCC patients while preserving the biological variation of the data. UMAP analysis showed clear improvements of data homogeneity upon batch correction. Notably, the renal cell cancer diagnosis sub-groups clustered according to their phenotype. This becomes clear as chRCC (malign) and OC (benign) cluster together and they are often pathologically indistinguishable due to similar cell origins.

Hierarchical clustering of MSI data of RCC tumor cores improves upon batch correction. Healthy, OC and chRCC cluster subtype-specific only upon correction, compared to uncorrected that exhibits no specific grouping (Figure 1b). Upon comBAT correction the total variance across all features is evenly distributed across the batches as compared to the uncorrected data, which contained outliers (Figure 1d).

comBAT and CCA exhibited overall best performance with a score improvement of 19.4% vs uncorrected data. comBAT works on the intensity/counts data, which can be beneficial in terms of downstream analysis since single feature are retained after correction. The other three methods are reduced by PCA before correction and therefore makes single feature analysis challenging (e.g., identifying discriminative peaks). In the case of unsupervised data analysis, e.g. UMAP or hierarchical clustering, the tested batch correction methods are valid because the output does not rely on individual features.

We found that comBAT correction mainly affects the MSI mass range above *m/z* 2500 and below *m/z* 1000 (Figure 1g-i), indicating that ion signals in the range m/z 1000-2500 are reliable and exhibit low variation. This agrees with the fact that most tryptic peptide masses fall in the range of *m/z* 1000-2500.

In addition to the four batch correction methods included here, several autoencoders for single cell omics data analysis are available in the Python programming language, such as scVI[18], scANVI[19] or Scanorama[20]. They may find use in MSI data analysis by using the MSI data formatting and encoding presented in our study. However, MSI data has a dense data structure that may be incompatible with the fact that existing autoencoders performs best for sparse data. Autoencoder based correction may therefore require adaption for MSI data.

All the included methods are available through R. Additional methods exist in Python code including neural networks that perform well for scRNA seq data.

## Conclusion

We introduce an MSI data structure that allows batch correction of large-scale mass spectrometry imaging data by methods adapted from the single cell omics fields. The underlying biological variation is conserved after batch correction with a score improvement of up to 19.4%. Batch correction allows for more robust and more accurate large-scale MSI studies of sample cohorts and is an important step towards wider applications of MSI in pharmacology and clinical research, including pathology.

## Acknowledgements

AAT was financed by a PhD fellowship from University of Southern Denmark (SDU), Department of Biochemistry and Molecular Biology. The authors thanks Maj Rabjerg, MD and Niels Marcussen, MD, OUH Dept. Pathology for providing samples. AAT thanks DANEMO, the Danish EMBL node, for financial support of a short-term visit to Theodore Alexandrov and his research group at the European Molecular Biology Laboratory, Heidelberg, Germany, to discuss data processing in MSI.

